# Rapid Visual Engagement in Neural Processing of Detailed Touch Interactions

**DOI:** 10.1101/2024.08.29.610402

**Authors:** Sophie Smit, Almudena Ramírez-Haro, Genevieve L. Quek, Manuel Varlet, Denise Moerel, Tijl Grootswagers

## Abstract

Touch perception is an inherently multisensory process in which vision plays an essential role. However, our understanding of how vision encodes sensory and emotional-affective aspects of observed touch, and the timing of these processes, is still limited. Here we address this gap by investigating the neural dynamics of visual touch observation by analysing electroencephalographic (EEG) data from participants viewing detailed hand interactions from the Validated Touch-Video Database. We examined how the brain encodes basic body cues such as hand orientation and viewing perspective, in addition to sensory aspects, including the type of touch (e.g., stroking vs. pressing; hand vs. object touch) and the object involved (e.g., knife, brush), as well as emotional-affective dimensions. Using multivariate decoding, information about body cues was present within 60 ms, with sensory and emotional details, including valence, arousal, and pain present around 130 ms, demonstrating efficient early visual processing. Threat was most clearly identified by approximately 265 ms, similarly involving visual regions, suggesting that such evaluations require slightly extended neural engagement. Our findings reveal that bottom-up, automatic visual processing is integral to complex tactile assessments, important for rapidly extracting both the personal relevance and the sensory and emotional dimensions of touch.

## Introduction

When perceiving touch, our brain extracts both visual body cues, such as the location of touch and whether it involves our body or another’s—indicated for instance by first or third-person perspectives—and sensory and affective information, like the pressure, texture, and emotional tone of the touch. It might seem that such diverse information requires complex high-level computations, yet evidence increasingly shows that recognition of social and affective cues in human interactions primarily relies on rapid, automatic, bottom-up visual processes, suggesting an evolutionarily adaptive mechanism (McMahon et al., 2023; McMahon and Isik, 2023; Pitcher and Ungerleider, 2021; Scholl and Gao, 2013). Recent theories propose that understanding social interactions, including their valence, goals, or intent, and possibly even the type of interaction, is fundamentally visual (McMahon and Isik, 2023). This does not imply mere low-level processing followed by cognitive interpretation, but rather suggests that our visual system contains advanced, abstract representations of social interactions. In observing social interactions that involve touch, such as two people hugging, aspects like valence and arousal are processed within 180 ms (Lee Masson and Isik, 2023). This highlights the brain’s capability to quickly discern the social-affective meaning of touch through feedforward visual processing (Lamme and Roelfsema, 2000), emphasising social touch interpretation as an integral aspect of visual perception. However, the visual encoding of other affective aspects like threat and pain perception in touch remain poorly understood, as do questions about the timing of such representations in relation to other tactile aspects, such as whether the touch involves an object or skin contact, and more nuanced sensory details, like the distinction between pressing or stroking. To address these questions, we investigate the neural dynamics of observed detailed touch interactions, examining how the brain encodes body cues, sensory qualities, and emotional-affective aspects.

Research on the neural basis of observed touch has often centred on the involvement of the somatosensory cortex (for reviews see Bufalari and Ionta, 2013; Gillmeister et al., 2017; Keysers and Gazzola, 2009; Peled-Avron and Woolley, 2022), an area of the brain traditionally linked to processing direct tactile inputs. These studies, driven by theories like those of mirror neurons (Gallese and Goldman, 1998; Keysers and Gazzola, 2009), propose that we internally replicate the sensory experiences of others as though the touch were happening to us, facilitating a form of ‘tactile empathy’ (Lamm et al., 2015; Marsh, 2018). For example, research indicates that observing touch enhances event-related potentials from direct tactile stimulation at both early sensory and later cognitive stages (Adler et al., 2016; Adler and Gillmeister, 2019; Bufalari et al., 2007; Galilee and McCleery, 2016; Martínez-Jauand et al., 2012; Pihko et al., 2010; Rigato et al., 2019b, 2019a; Smit et al., 2023; Streltsova and McCleery, 2014), suggesting that the somatosensory cortex simulates touch observed in others at various processing levels. However, others challenge this, suggesting that understanding others through touch observation can be achieved purely via visual processing, without the need for simulated experiences (de Vignemont, 2017; Hickok, 2014). This debate raises critical questions: Can the brain extract sensory and emotional details of touch purely through visual pathways, without involving the somatosensory cortex or other regions? Previous research has largely focused on these simulation processes through somatosensory regions, leaving a significant gap in our understanding of how purely visual processes encode the detailed sensory and emotional-affective aspects of perceived touch.

Here we test the neural dynamics of visual touch perception by using high-temporal resolution electroencephalography (EEG) to analyse brain responses as participants viewed detailed tactile hand interactions from the Validated Touch-Video Database (Smit and Rich, 2023).

Employing multivariate whole-brain decoding, we captured the rapid encoding of sensory and emotional details, such as the object involved and aspects like valence and arousal, within 130 ms, indicating swift initial visual processing. Additionally, we investigated for the first time the affective dimensions of pain and threat in touch contexts, demonstrating that these aspects are also rapidly detected within 135 and 285ms respectively, involving visual regions. Extending beyond prior studies focused on broader social contexts (Lee Masson and Isik, 2023; McMahon and Isik, 2023), our results emphasise the crucial role of rapid feedforward visual pathways in detecting sensory features and emotional salience in focused touch interactions, which are critical for quick, context-specific reactions.

## Methods

### Data availability

All raw data, stimulus presentation and analysis scripts, and video stimuli are available in BIDS format on OpenNeuro (doi:10.18112/openneuro.ds005439.v1.0.0).

### Ethics statement

This study was approved by the Western Sydney University ethics committee (project number: 15644) and informed written consent was obtained from all participants.

### Stimuli & video rating procedure

We used a set of stimuli previously developed and validated (The Validated Touch-Video Database: Smit and Rich, 2023). This original set includes 90 videos of varying lengths that depict various types of tactile interactions with a left hand shown from a first-person perspective (Fig. 1B). The videos differ along several dimensions, including arousal, perceived threat, hedonic qualities (neutral, pleasant, unpleasant, or painful), type of touch (e.g., stroking, pressing, stabbing), and whether the touch involved another hand or an object (such as a brush, hammer, or knife). For the current study, we standardised all video lengths to 600 milliseconds, centred on the touch event, comprising 15 frames at a rate of 25 frames per second. The videos were 256 pixels in width and 144 pixels in height presented at approximately 60 cm from the screen (6.2° visual angle). To explore different perspectives, we presented the videos in four orientations achieved by horizontal, vertical, or combined flips, illustrating touch to either the left or right hand from self-oriented or other-oriented viewpoints, totalling 360 stimuli.

**Figure 1.**
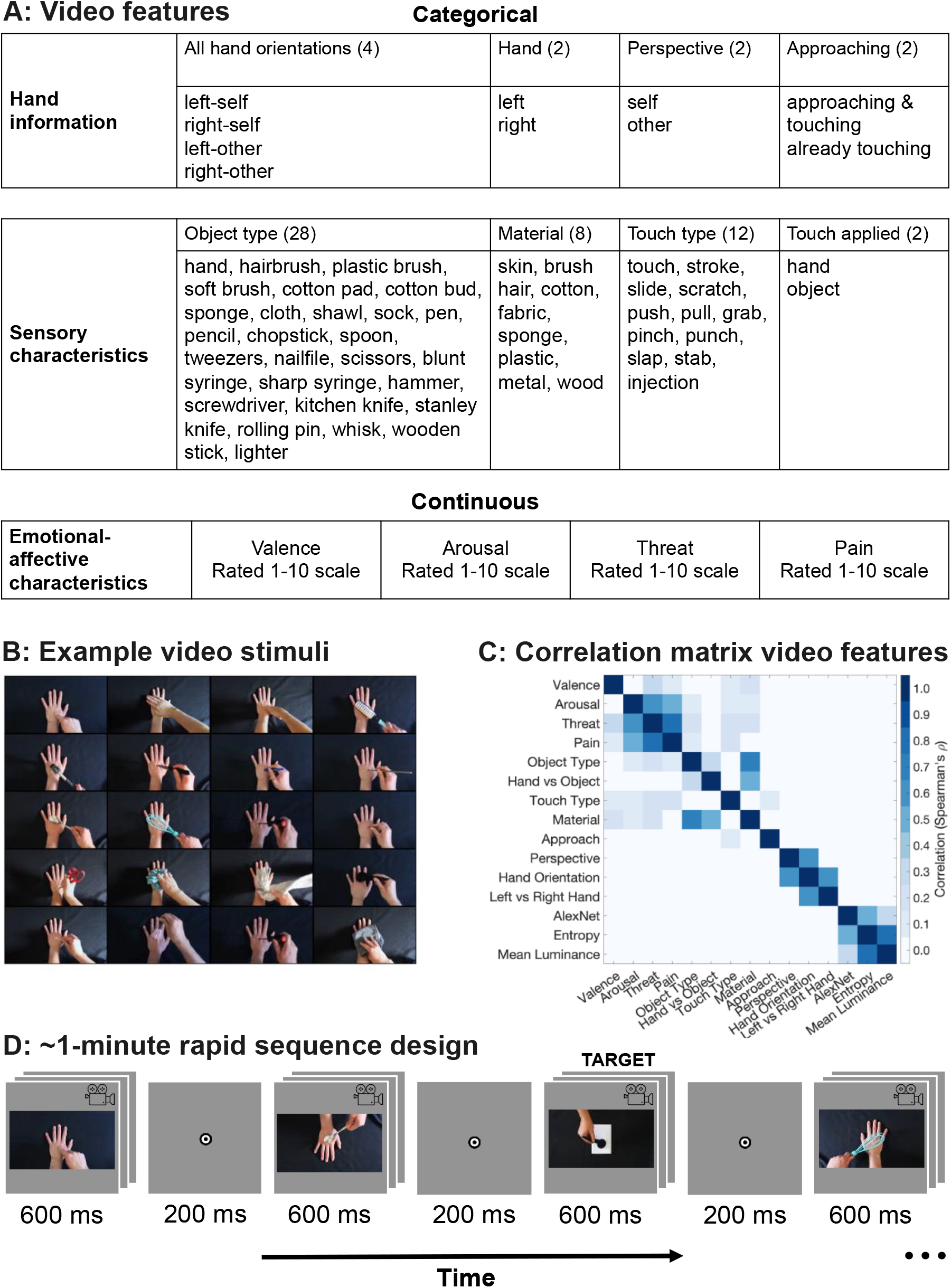
Experimental Stimuli and Design. A) Table of features used in the regression and decoding analysis, showing the categorical and continuous characteristics extracted from each video. B) Middle frames from a video subset, presented in four orientations during the experiment to depict left/right hand and self/other perspectives. C) The overlap of information between the different touch features related to the videos was assessed by creating an RDM for each feature and correlating the models with one another. D) EEG experiment design with an example sequence. Participants initiated by pressing the space bar; videos appeared for 600 ms with 200 ms intervals. The task involved counting targets trials where a white object, rather than a hand, was touched. Each sequence duration ranged from 54 to 78 seconds.

To confirm that our modifications (shorter duration, smaller presentation size, and different orientations) did not impact the perceived attributes of the touch compared to the original videos that were previously rated, we recruited a separate cohort of participants to evaluate the modified clips using the same criteria as the original dataset (these participants were independent from the participants in the EEG task). There were 82 participants (68 female, 14 male, mean age 26.3, age range 18-57 years) for the online video rating task. Participants were recruited through Western Sydney University and received course credit or payment. We administered four distinct questionnaires via Qualtrics, each showing the abbreviated 90 videos in only one of the four orientations (e.g., a participant would only see touches to a right hand in a first-person perspective). Participants were restricted to one questionnaire each, ensuring separate ratings, with 20 or 21 participants contributing ratings for each questionnaire. We included the same four questions from the original database study: (1) How would you categorise the touch in this video? (single answer: neutral, pleasant, unpleasant, painful); (2) How [pleasant/unpleasant/painful] (based on the previous answer) was the touch?; (3) How threatening was the touch?; and (4) How arousing was this video? (arousing in terms of feeling, emotion, or response). The last three questions were rated on a scale from 1 (not at all) to 10 (extremely). The results indicate strong correlations with old ratings (see supplementary Table 1). We use the arousal, threat, and pain ratings from the original video database in our analyses, as described below. Valence, however, was not as clearly captured by the original ratings, as participants categorised videos as neutral, pleasant, unpleasant or painful and then only rated that dimension on a scale from 1-10. To generate a single valence score for each video, we used the percentage of participants who categorised the video as either pleasant or unpleasant and then applied Principal Component Analysis (PCA) to these percentages. We extracted the first principal component to create a unified valence score.

### EEG experiment

We recruited 32 participants for the EEG experiment with (25 female, 7 male, mean age 28.3 years, age range 18–51 years, 19 right-handed and 3 left-handed). Participants were recruited through Western Sydney University and received course credit or payment. We used a rapid sequence design (e.g., Grootswagers et al., 2022, 2019) where each participant was presented with 32 sequences including 90 different videos presented in four orientations (360 video stimuli total) showing touch to a hand presented in a fully random and counterbalanced order. To maintain participant attention, additional target stimuli, which involved touch interactions with a white object rather than a hand, were interspersed randomly among the touch videos and were excluded from the analysis (Fig. 1B). Each sequence contained 1 to 9 target stimuli, with a minimum of 12 non-target stimuli between consecutive targets. Participants were tasked with counting these targets and reporting their count at the end of the sequence via the top row numbers on the keyboard, receiving immediate feedback on their accuracy. Participants achieved an average accuracy of 81.93% (SD = 18.89%) on this target-detection task, demonstrating their high engagement. Each sequence ranged in duration from 54 to 78 seconds, depending on the number of target trials. The videos, each 600 ms in duration, were separated by a 200 ms inter-video gap and displayed against a light grey background on a 24-inch ViewPixx monitor. Participants were instructed to take breaks as needed in between sequences and self-initiated the next sequence though a keypress. The EEG task included a total of 2880 non-target trials (each video presented eight times) alongside a variable number of target trials, going for approximately 55 minutes including breaks. The experiment was conducted using Python and PsychoPy software version 2023.3.1 (Peirce et al., 2019).

Additionally, we administered four short questionnaires at the end of the experiment, which required an extra 15 minutes to complete. These are not included in the current analyses.

### EEG recordings and preprocessing

EEG data were continuously recorded using a 64-channel BioSemi Active-Two electrode system at a sampling rate of 2048 Hz (BioSemi, Amsterdam, The Netherlands). Voltage offsets were maintained below 20 mV. The electrode placement adhered to the 10/20 international standard (Jasper, 1958; Oostenveld and Praamstra, 2001), offline preprocessing was performed using the Python MNE toolbox version 1.7 (Gramfort et al., 2014). We re-referenced the data to a common average followed by band-pass filtering using a 0.1 Hz high-pass and a 100 Hz low-pass filter to remove slow drifts and high-frequency artifacts, including muscle noise. The data were then downsampled to 200 Hz to reduce the data size and computational load. Baseline correction was applied using a window from -100 to 0 ms before stimulus onset. Temporal smoothing was accomplished with a moving average filter over 10 samples (40 ms), to reduce short-term fluctuations. Epochs were extracted from -100 to 800 ms relative to each stimulus onset, with no additional preprocessing. All voltages from each channel at every time point were used for subsequent analyses.

### Decoding analysis

To analyse the neural processing of both continuous and categorical aspects of observed visual touch in the EEG data (Fig. 1A), we employed a combination of time-resolved regression and classification analyses (Grootswagers et al., 2017). Continuous aspects, including subjective ratings of threat, arousal, valence, and pain from the Validated Touch-Video Database (Smit and Rich, 2023: rated on a scale from 1 to 10), were predicted using regression. Categorical aspects included all four hand orientations (i.e., hand x perspective), hand (i.e., left vs. right), and perspective (i.e., self vs. other), whether a hand was coming in to touch the other hand or whether the touch was already occurring from the start of the video, object type (e.g., hand, brush), material (e.g., metal, wood; or skin when touched by a hand), touch type (e.g., touch, stroke), and whether the touch was applied by another hand or an object. These categories were all manually coded for this study and decoded using classification. This approach allowed us to assess how accurately the EEG data, channel voltages recorded from 64 electrodes, could predict the subjective ratings taken from the video database (note the videos were rated by a large and separate sample of participants: Smit and Rich, 2023) and distinguish between different categories of touch.

For both the classification and regression analysis, we employed a sliding estimator over time, training and testing on the same timepoint, in combination with leave-one-sequence-out cross-validation. In each iteration of the cross-validation, we trained our models on 31 sequences and tested on the remaining sequence, repeating this process 32 times. This approach ensured that each sequence was used as a test set once. For the classification and regression, we first standardised all input features by transforming the data to have a mean of zero and a standard deviation of one, preventing features with larger scales from disproportionately influencing the model. For the regression analysis, we used ridge regression with an alpha parameter of 0.5, which helps to control for overfitting by adding a penalty to the model coefficients. The regression model was trained on EEG data to predict continuous emotional dimensions (valence, arousal, threat, and pain) rated on a 1-10 scale for each video in a separate study (Smit and Rich, 2023). Prediction performance was assessed by correlating the model’s cross-validated ratings predicted from the EEG data with the subjective video ratings, with this correlation compared to a chance level of zero. In the classification analysis, we applied regularised Linear Discriminant Analysis (LDA) to classify patterns of neural activity associated with the different categorical features. The performance of the classification model was evaluated using balanced accuracy, which accounts for class imbalances by averaging the accuracy across all classes, ensuring that the classification performance is not biased towards more frequent classes. The balanced accuracy was then compared to a chance level, which was determined based on the number of possible labels (e.g., with 10 labels, the chance level would be 10%). To assess the spatial contributions to classification and regression performance, we conducted a channel searchlight analysis at nine specified single time points (−50, 50, 150 etc. to 750 ms), calculating correlation scores and classification accuracies for each EEG channel, including their adjacent channels, instead of the whole brain as in the main analyses. These analyses were conducted in Python using the MNE toolbox (Gramfort et al., 2014).

To account for potential confounding visual effects, we controlled for key visual characteristics by extracting and regressing out three distinct visual features from the EEG data: entropy, mean luminance, and AlexNet features. Entropy and mean luminance were estimated from the first frame of each video. Entropy was calculated as a measure of the randomness or complexity within the grayscale image, reflecting the variability in pixel intensities. Mean luminance was computed as the average brightness of the pixels in the coloured image, providing an effective measure of the overall lightness of the frame. The choice of the first frame is justified by its role in setting the visual tone and brightness for the video. For the AlexNet features, we used the coloured middle frame of each video, inputting it into AlexNet layer 11 and converting the output into a one-dimensional feature vector using PCA. The middle frame was selected to capture the typical content of the video, which likely reflects the central theme or action, making it a good choice for extracting deep neural network features. The three visual models were created in MATLAB and the results were loaded into Python for subsequent analyses.

Entropy and mean luminance were calculated using standard image processing libraries, while the AlexNet features were derived by applying a pre-trained deep learning model (Krizhevsky et al., 2012). Each of these visual features—entropy, mean luminance, and the AlexNet PCA component—was treated as a separate model in our regression analysis. We regressed out these features from the EEG signal by fitting a linear regression model at each channel and time point within each fold of the cross-validation process. The residuals from this regression were then used for fitting the classification and regression models, ensuring that the EEG data reflected neural responses to touch perception rather than variance due to visual input.

To assess the association between various features in our decoding analysis, we used dissimilarity matrices (Kriegeskorte and Kievit, 2013). These were generated from perceptual ratings from the Validated Touch-Video Database (Smit and Rich, 2023) for continuous features, manual coding results for categorical features, and results from the visual models AlexNet, entropy, and mean luminance. Dissimilarity matrices are particularly useful as they allow for the comparison of both continuous and categorical data on a common scale, facilitating a comprehensive analysis across different data types (Kriegeskorte and Kievit, 2013). We employed Spearman’s rank correlation to evaluate the relationships among these models, uncovering how different characteristics correlate with one another (Fig 1C).

### Statistical inference

We used Bayes factors to evaluate the evidence for both the null (chance decoding) and alternative (above-chance decoding) hypotheses (Dienes, 2011; Kass and Raftery, 1995; Morey et al., 2016; Rouder et al., 2009). Bayes factors were computed using the ‘BayesFactor’ package from R (Morey et al., 2018), integrated into Python via the rpy2 interface. We applied a half-Cauchy prior for the alternative hypothesis to capture above-chance effects. The prior was centred around chance with the default width *r*=0.707 (Jeffreys and Jeffreys, 1998; Rouder et al., 2009; Wetzels et al., 2011). Based on previous recommendations (Teichmann et al., 2022), we omitted the interval from d = 0 to d = 0.5 from the prior to disregard effect sizes considered too small to be relevant. However, this approach was not used for the topoplots, where we anticipated smaller effect sizes due to fewer sensors capturing more localised patterns. Bayes factors quantify the strength of evidence, with a Bayes factor greater than 1 indicating support for the alternative hypothesis, and less than 1 supporting the null (Jeffreys and Jeffreys, 1998). A Bayes factor of 6, for example, suggests that the evidence favouring the alternative hypothesis is six times stronger than that for the null, although we avoid strict cut-offs to maintain a focus on the evidence continuum rather than binary outcomes (Dienes, 2011; Kass and Raftery, 1995; Morey et al., 2016; Rouder et al., 2009).

## Results

We explored the temporal dynamics of neural responses to visually perceived touch. We recorded EEG data while participants watched brief and detailed touch interaction videos, adjusted from the Validated Touch-Video Database (Smit and Rich, 2023). This study aims to address a gap in our knowledge of visual touch perception by examining how the brain processes visual body cues, such as viewing perspective, alongside more complex sensory and emotional-affective interpretations of touch.

We decoded categorical features related to the observed touch at each time-point to determine when this information appears in the EEG responses. We first focused on the viewing perspective and movement dynamics (Fig. 2). Our findings reveal that hand orientation is decoded very swiftly within the initial 60 ms and peaking around 125 ms, including distinctions of hand (left vs. right) and perspective (self vs. other). A distribution of channels across occipital, central, and frontal regions contributed to the decoding of these aspects. We did not find evidence for above-chance decoding of the more intricate interaction dynamics comparing whether one hand is approaching another or if the touch is already occurring from the first frame. These results indicate that visual information about the body part touched and the likely recipient (self or other) is processed rapidly via the visual pathway.

**Figure 2.**
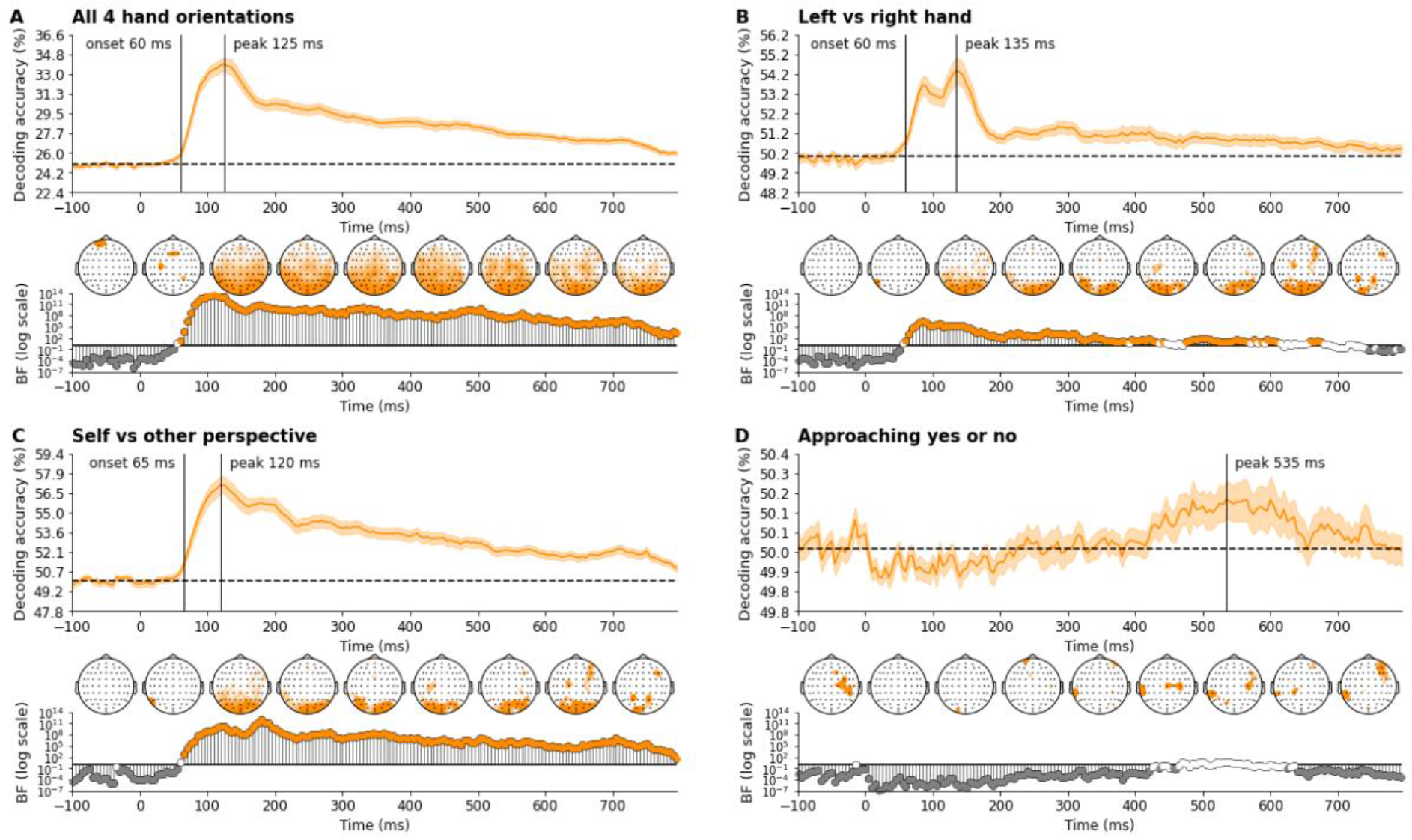
Time course of decoding accuracies of hand information show rapid discrimination between hand orientations and perspectives, with a notably later response for approaching movements. These plots illustrate the time-varying decoding accuracies for A) all four hand orientations, B) left versus right hand, C) self versus other perspective, and D) the approach of one hand towards another versus an already touching hand. Stimulus onset is at 0 ms. Theoretical chance levels are set at 25% for the four hand orientations and 50% for binary distinctions (e.g., left vs. right hand, self vs. other perspective, approaching yes or no). Shaded areas around the plot lines represent the 95% confidence intervals across participants (N = 30). Below the plots, Bayes factors are displayed on a logarithmic scale. Bayes factors below 1/6, depicted in grey, signify strong evidence for the null hypothesis, while those above 6, shown in colour, suggest strong support for the alternative hypothesis. Bayes factors between 1/6 and 6 are indicated in white. Vertical lines show the onset and peak of sustained above-chance decoding. Additionally, time-varying topographies derived from the channel-searchlight analysis are presented in colour, showing Bayes factors ≥ 3 for individual time points ranging from -50 ms to 750 ms at 100 ms intervals.

Next, we examined the sensory characteristics of the observed touch (Fig. 3). The type of object and related material attributes, which varied in texture and other properties (e.g., metal may be perceived as cold and a cloth as warm), were both decoded rapidly within 150 ms. The ability to distinguish between a touch applied by a hand, involving direct skin contact, or with an object, emerged slightly later, by about 200 ms. Information about these sensory aspects was sustained for some time, with peaks occuring around 200-300ms, indicating the time when most information was present. Processing of sensory aspects appeared to recruit visual brain regions, moving more central and then frontal/temporal over time, especially for the material involved in the touch. The type of touch (e.g., stroking versus pressing) was not clearly decodable, suggesting this may be a less distinct feature in the videos. These findings highlight representations of object-related sensory information in the brain’s initial visual processing stages.

**Figure 3.**
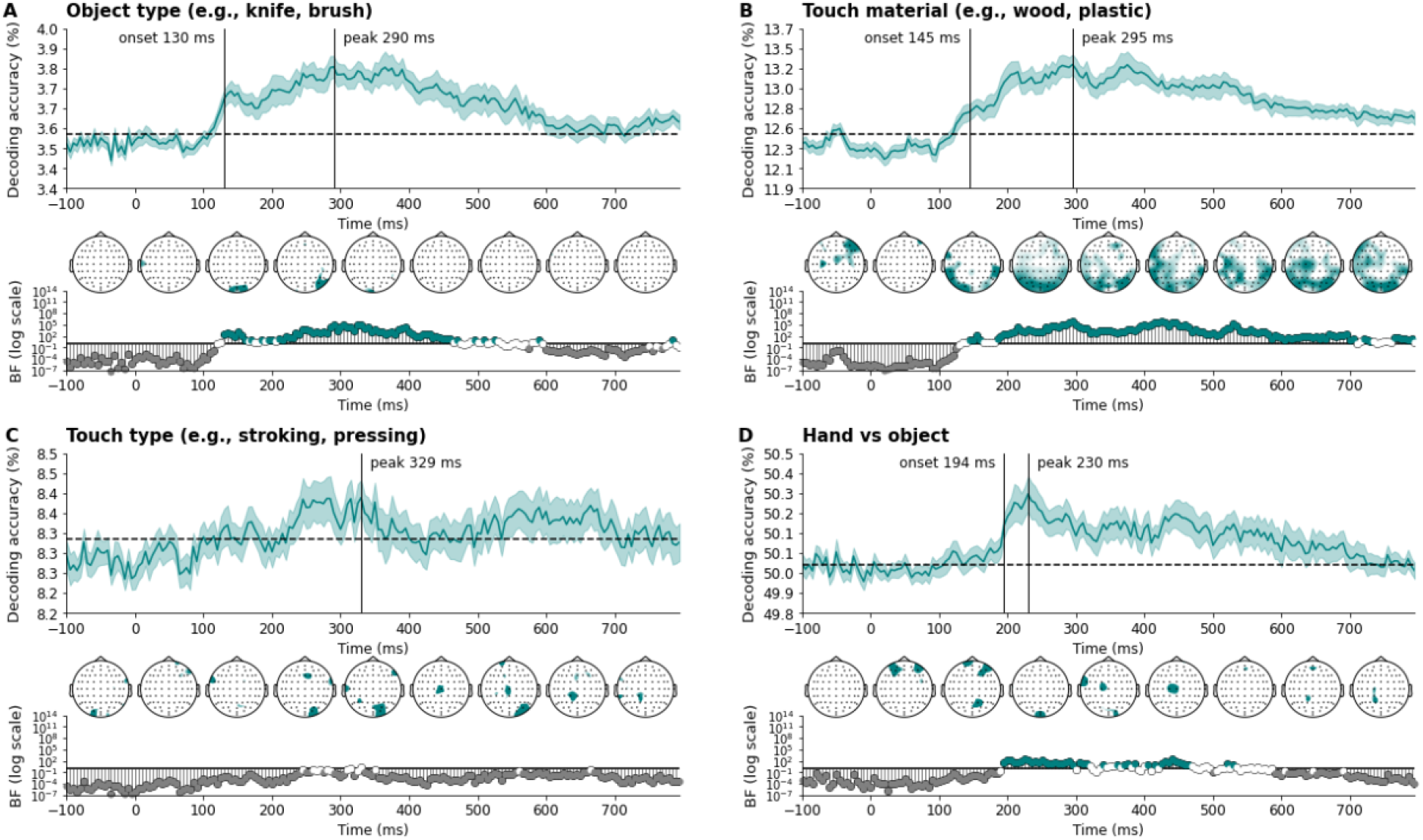
Time course of decoding accuracies of sensory characteristics show rapid discrimination of object, material, and whether the touch was applied by a hand or an object. These plots illustrate the time-varying decoding accuracies for A) object type, B) the material involved in the touch (note that contact between hands is included as a label), C) touch type, and D) touch applied by a hand vs. an object. Stimulus onset is at 0 ms. Theoretical chance levels are set at 3.6% for object type, 12.5% for material, 8.3% for touch type, and 50% for hand vs. object. Shaded areas around the plot lines represent the 95% confidence intervals across participants (N = 30). Below the plots, Bayes factors are displayed on a logarithmic scale. Bayes factors below 1/6, depicted in grey, signify strong evidence for the null hypothesis, while those above 6, shown in colour, suggest strong support for the alternative hypothesis. Bayes factors between 1/6 and 6 are indicated in white. Vertical lines show the onset and peak of sustained above-chance decoding. Additionally, time-varying topographies derived from the channel-searchlight analysis are presented in colour, showing Bayes factors ≥ 3 for individual time points ranging from -50 ms to 750 ms at 100 ms intervals.

Last, we looked at the emotional-affective aspects of the observed touch (Fig. 4). Valence and arousal showed a very similar time course, being present in the neural data from 130 ms with peaks around 300ms. This suggests an early neural encoding of these affective dimensions, which likely reflects rapid processing of emotional salience. Especially valence was clearly present and sustained in the data, showing initial activation of visual regions and progressively moving more central. Information about the level of pain was present in the data at a similar time, from approximately 135 ms, with the perception of threat detected at 265 ms, indicating a more delayed response that may involve higher cognitive integration to evaluate potential danger. Though the decoding of threat occurred somewhat later in time, with pain and threat both peaking around 380-400 ms, the spatial activations still suggest involvement from visual regions. This progression illustrates that while more basic emotional responses to touch are processed quickly, and via the initial visual processing pathway, more complex evaluations about potential harm involve slightly longer neural processing times and potentially deeper cognitive involvement.

**Figure 4.**
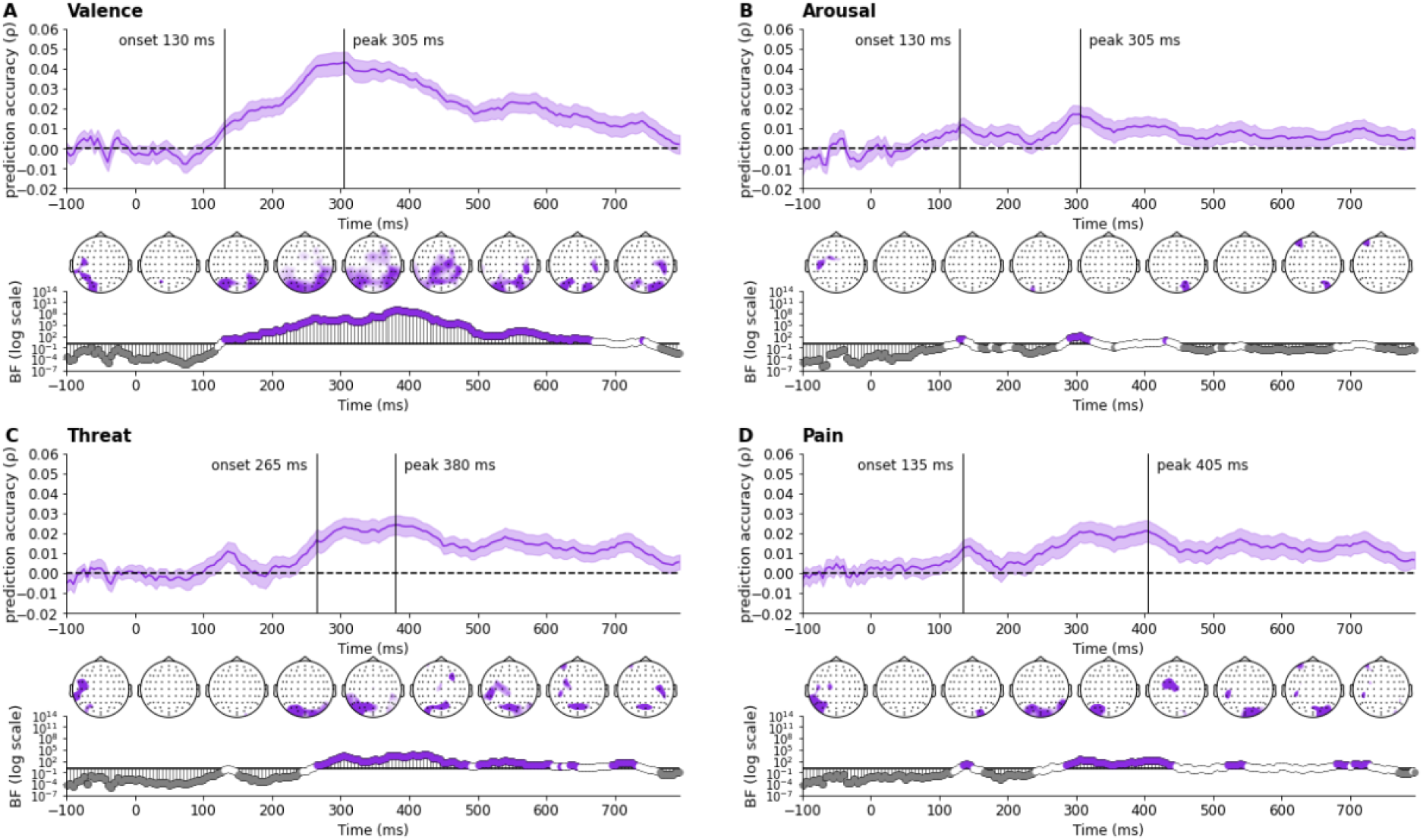
Time course of decoding accuracies of emotional-affective characteristics show rapid discrimination of basic emotional responses valence and arousal, with a slightly later response for more complex evaluations of threat and pain. These plots illustrate the prediction accuracy for A) valence, B) arousal, D) threat, and D) pain. Stimulus onset is at 0 ms. Shaded areas around the plot lines represent the 95% confidence intervals across participants (N = 30). Below the plots, Bayes factors are displayed on a logarithmic scale. Bayes factors below 1/6, depicted in grey, signify strong evidence for the null hypothesis, while those above 6, shown in colour, suggest strong support for the alternative hypothesis. Bayes factors between 1/6 and 6 are indicated in white. Vertical lines show the onset and peak of sustained above-chance decoding. Additionally, time-varying topographies derived from the channel-searchlight analysis are presented in colour, showing Bayes factors ≥ 3 for individual time points ranging from -50 ms to 750 ms at 100 ms intervals.

## Discussion

We examined the neural dynamics of visually perceived touch using EEG. Participants viewed 90 brief video clips, adjusted from the Validated Touch-Video Database (Smit and Rich, 2023), displaying a range of touch interactions, and presented here as either a left or right hand from a self or other specifying perspective. These videos, rated on various dimensions, were selected to probe how the brain interprets touch from visual body cues to more complex aspects. Using multivariate decoding, we tested the neural representation of key touch dimensions such as hand orientation, touch quality, and emotional-affective implications.

Our study, focusing on close-up views of hands involved in various touch interactions, builds on prior research that predominantly examined the neural dynamics of touch observation within social contexts (Lee Masson et al., 2020, 2018; Lee Masson and Isik, 2023; Peled-Avron et al., 2016; Peled-Avron and Shamay-Tsoory, 2017; Schirmer and McGlone, 2019). By emphasising the specific details of touch rather than broader social interactions, we have conducted an in-depth exploration of how the brain processes different touch-related elements, revealing that in addition to body cues, sensory and complex emotional aspects such as object type, material, valence, and arousal are also rapidly decoded by the brain’s initial visual pathways.

Importantly, we carefully controlled for low-level visual information in the data to ensure that any observed effects are attributed to meaningful aspects of observed touch, not to structural variations in the video stimuli. The encoding of relevant emotional information in visual regions—observed as early as 130 ms and peaking around 300 ms for valence and arousal— parallels findings from observed social touch interactions, where these aspects are processed within a similar timeframe (170 ms and 180 ms respectively, Lee Masson and Isik, 2023). This consistency across contexts emphasises the critical role of initial visual pathways in recognising emotional salience, with activation in visually-selective areas occurring prior to engagement of the somatosensory cortex (Lee Masson and Isik, 2023). Our findings align with broader research indicating that the processing of complex social-emotional information, including the perception of valence and arousal in social interactions (McMahon and Isik, 2023), and emotional images (Grootswagers et al., 2020), is managed automatically and in a bottom-up fashion by visually-selective brain regions. These capabilities of the visual system are essential for quick and effective responses to tactile events, demonstrating its essential role in interpreting multifaceted touch interactions.

We extended our investigation to the affective dimensions of threat and pain. We observed that pain perception was similarly decoded around 135 ms with later, more sustained activations around 300-400 ms. This time course aligns with prior research on visual depictions of hands in painful scenarios, which identified ERP modulations differentiating painful from neutral stimuli at 140 ms and again at 380 ms, with early amplitudes (140-180 ms) correlating with subjective assessments of observed pain and personal discomfort (Fan and Han, 2008). Although previous research has investigated the time course of threat perception in other contexts like facial expressions, showing modulations between 120 and 280 ms and later around 400-500 ms (Schupp et al., 2004; Williams et al., 2006), little is known about the temporal dynamics of threat perception in the context of observed touch. In our study, the perception of threat emerged most clearly around 265 ms and peaked at 380 ms, suggesting a somewhat slower response compared to the other emotional touch aspects, likely requiring greater cognitive effort to assess potential danger. Despite this delayed timing, however, the similar involvement of visual regions indicates that both immediate and complex emotional responses to touch are integrated within the visual processing system.

Analysing spatial patterns over time, we found that the decoding of sensory and emotional aspects, especially valence and material properties, begins in visual regions and quickly extends to central and frontal areas, and for material properties, progresses into temporal regions. This transition from visual to central regions may potentially reflect a simulation of the sensory properties of observed touch within the observer’s somatosensory cortex (Gallese et al., 2004; Keysers and Gazzola, 2009; Lee Masson and Isik, 2023; Peled-Avron and Woolley, 2022). In addition to valence, the brain’s assessment of materials involved in touch presumably includes both physical properties and emotional or social implications, such as the comfort of soft fabrics versus the discomfort of rough surfaces, and the intimacy of skin-to-skin versus object-to-skin contact. Processing these dimensions of touch would likely involve temporal regions associated with extracting body postures and interactions, including emotional content, such as the extrastriate body area, fusiform body area, and posterior superior temporal sulcus (Keysers and Gazzola, 2006; Peelen et al., 2007; Peelen and Downing, 2007).

Additionally, regions critical for object property recognition, such as the lateral occipital complex and the fusiform gyrus, likely play a role (Kanwisher et al., 1996; Malach et al., 1995). While prior research on focused touch interactions has primarily emphasised the role of the observer’s somatosensory cortex in creating a vicarious sensory experience (Adler et al., 2016; Adler and Gillmeister, 2019; Bufalari et al., 2007; Galilee and McCleery, 2016; Martínez-Jauand et al., 2012; Pihko et al., 2010; Rigato et al., 2019b, 2019a; Smit et al., 2023; Streltsova and McCleery, 2014), our study reveals an essential involvement of visual areas in rapidly processing touch-related information, which then propogates to more central, temporal, and frontal regions.

Future research could delve deeper into how individual differences in traits like empathy and vicarious touch experiences influence the neural processing of visually perceived touch. For instance, individuals with heightened vicarious touch experiences, who physically feel sensations when observing others being touched (Gillmeister et al., 2017; Smit et al., 2024), may process observed touch stimuli differently compared to those without such experiences. These individuals might exhibit temporal variations in the encoding of tactile information (Smit et al., 2023), with potentially less distinction between self and other perspectives due to a diminished clarity in self-other differentiation (Banissy et al., 2009; Ward and Banissy, 2015).

Another extension of this research is to investigate the neural dynamics of purely sensory touch interactions devoid of any social context. While our study focused on close-up touches to isolate touch details, it still inherently involved a social element due to the interaction between two hands. Future studies could eliminate this aspect by showing mechanical or object-mediated touch interactions, allowing for a clearer examination of sensory processing without the influence of social context.

In conclusion, our findings reveal the significant role rapid and automatic visual processing plays in extracting the sensory and emotional characteristics of perceived touch. This has broader implications for developing more effective and nuanced models of sensory processing, and for applications in areas that focus on multisensory integration, such as neuroprosthetics where visual feedback could enhance the perception of touch.

## Acknowledgements

This research was supported by Australian Research Council grants DP220103047 and DE230100380.

