## Supplementary Materials for "Rapid Visual Engagement in Neural Processing of Detailed Touch Interactions"

**Supplementary Table 1:** Correlations between ratings for the adapted videos used in this study and the original videos from the Validated Touch-Video Database.

| Video Orientation | Touch Dimension | Correlation (r) | Bayes Factor |
|---|---|---|---|
| Normal | Neutral | 0.88 | 59.69 |
| Normal | Pleasant | 0.95 | 92.41 |
| Normal | Unpleasant | 0.82 | 44.51 |
| Normal | Painful | 0.93 | 82.61 |
| Normal | Threat | 0.97 | 109.24 |
| Normal | Arousal | 0.88 | 61.02 |
| Horizontal Flip | Neutral | 0.86 | 54.11 |
| Horizontal Flip | Pleasant | 0.94 | 86.41 |
| Horizontal Flip | Unpleasant | 0.82 | 45.37 |
| Horizontal Flip | Painful | 0.91 | 70.01 |
| Horizontal Flip | Threat | 0.95 | 94.6 |
| Horizontal Flip | Arousal | 0.86 | 53.16 |
| Vertical Flip | Neutral | 0.85 | 51.08 |
| Vertical Flip | Pleasant | 0.93 | 82.16 |
| Vertical Flip | Unpleasant | 0.84 | 49.46 |
| Vertical Flip | Painful | 0.9 | 67.54 |
| Vertical Flip | Threat | 0.95 | 94.58 |
| Vertical Flip | Arousal | 0.87 | 55.73 |
| Hor & Vert Flip | Neutral | 0.83 | 45.8 |
| Hor & Vert Flip | Pleasant | 0.92 | 76.84 |
| Hor & Vert Flip | Unpleasant | 0.85 | 51.11 |
| Hor & Vert Flip | Painful | 0.9 | 65.38 |
| Hor & Vert Flip | Threat | 0.95 | 96.65 |
| Hor & Vert Flip | Arousal | 0.88 | 58.53 |
